# High-confidence Coding and Noncoding Transcriptome Maps

**DOI:** 10.1101/109363

**Authors:** Bo-Hyun You, Sang-Ho Yoon, Jin-Wu Nam

## Abstract

The advent of high-throughput RNA-sequencing (RNA-seq) has led to the discovery of unprecedentedly immense transcriptomes encoded by eukaryotic genomes. However, the transcriptome maps are still incomplete partly because they were mostly reconstructed based on RNA-seq reads that lack their orientations (known as unstranded reads) and certain boundary information. Methods to expand the usability of unstranded RNA-seq data by predetermining the orientation of the reads and precisely determining the boundaries of assembled transcripts could significantly benefit the quality of the resulting transcriptome maps. Here, we present a high-performing transcriptome assembly pipeline, called CAFE, that significantly improves the original assemblies, respectively assembled with stranded and/or unstranded RNA-seq data, by orienting unstranded reads using the maximum likelihood estimation and by integrating information about transcription start sites and cleavage and polyadenylation sites. Applying large-scale transcriptomic data comprising ninety-nine billion RNAs-seq reads from the ENCODE, human BodyMap projects, The Cancer Genome Atlas, and GTEx, CAFE enabled us to predict the directions of about eighty-nine billion unstranded reads, which led to the construction of more accurate transcriptome maps, comparable to the manually curated map, and a comprehensive lncRNA catalogue that includes thousands of novel lncRNAs. Our pipeline should not only help to build comprehensive, precise transcriptome maps from complex genomes but also to expand the universe of non-coding genomes.

## Introduction

Comprehensive transcriptome maps enhance understanding of gene expression regulation in both coding and noncoding genomic regions (Wang et al. 2009; Martin and Wang 2011). For the last decade, transcriptome-wide analysis, including genome-wide tiling array applications (Bertone et al. 2004; Cheng et al. 2005; Kapranov et al. 2005) and high-throughput RNA sequencing (RNA-seq) (Cloonan et al. 2008; Mortazavi et al. 2008; Nagalakshmi et al. 2008; Salehi-Ashtiani et al. 2008; Wilhelm et al. 2008), have unveiled pervasive transcription signals across genomes from both unicellular and complex multicellular organisms (Jacquier 2009; Croucher and Thomson 2010; Gerstein et al. 2010; Djebali et al. 2012; Harrow et al. 2012; Brown et al. 2014; Fort et al. 2014; Martin et al. 2014; Maudhoo et al. 2014; Moreton et al. 2014). Recently, large-scale RNA-seq data from the ENCODE project were used to characterize highly complex, overlapping transcription units on both strands, revealing that more than 60% of the human genome is reproducibly transcribed in at least two different cell types (Djebali et al. 2012; Harrow et al. 2012). Intriguingly, a significant portion of these extensive transcription signals, mostly from intergenic regions, turned out to be unannotated. To identify the unannotated transcriptome, gene annotation projects, such as GENCODE (Harrow et al. 2012), Human BodyMap (Cabili et al. 2011), and MiTranscriptome (Iyer et al. 2015), have massively reconstructed whole transcriptomes by assembling large-scale RNA-seq data and have characterized transcriptome-wide noncoding RNAs (ncRNAs). The majority of RNAs in the noncoding transcriptome were long ncRNAs (lncRNAs), such as repeat-associated ncRNAs, enhancer-associated ncRNAs, long intervening ncRNAs (lincRNAs), antisense ncRNAs, and so on, (Ulitsky et al. 2011; Derrien et al. 2012; Nam and Bartel 2012; Pauli et al. 2012; Young et al. 2012; Hangauer et al. 2013; Luo et al. 2013; Brown et al. 2014; Iyer et al. 2015).

Unknown transcripts can be identified via assembly of RNA-seq data by two approaches: the genome-guided approach (known as reference-based assembly) (Yassour et al. 2009; Guttman et al. 2010; Trapnell et al. 2010; Boley et al. 2014; Mangul et al. 2014; Maretty et al. 2014) and the *de novo* approach (Martin et al. 2010; Grabherr et al. 2011; Schulz et al. 2012; Safikhani et al. 2013; Xie et al. 2014; Chang et al. 2015; Pertea et al. 2015; Tjaden 2015). Because the *de novo* approach assembles RNA-seq reads without a guide genome, it generally requires RNA-seq data with strand information (called stranded RNA-seq data). However, for the reference-based approach, the stranded RNA-seq data had been regarded as dispensable because the sense-orientation of some reads spanning exon junctions could be predicted based on the splicing signal. For that reason, the Human ENCODE project (de Souza 2012), the modENCODE project (Gerstein et al. 2010; mod et al. 2010), the Human BodyMap project (Cabili et al. 2011), the Genotype-Tissue Expression (GTEx) project (http://www.gtexportal.org) (Consortium 2013), the human protein atlas (Uhlen et al. 2010; Uhlen et al. 2015), and the Cancer Genome Atlas (TCGA) (Ciriello et al. 2013; Kandoth et al. 2013) consortium produced large-scale unstranded RNA-seq data that lack strand information; genome-wide gene annotation projects have proceeded using these data. For instance, the MiTranscriptome project re-used 6,810 publicly available unstranded RNA-seq data from ENCODE, TCGA, and other studies to reconstruct a comprehensive map of the noncoding transcriptome (Iyer et al. 2015). Despite such applications, transcriptome assembly using unstranded RNA-seq data often results in erroneous transcript models, including chimeras, particularly when there are convergent, divergent, or antisense overlaps between two genes (Garber et al. 2011; Martin and Wang 2011; Nam and Bartel 2012). Stranded RNA-seq data can benefit reference-based assembly in those genomic regions (Martin and Wang 2011). Nevertheless, the re-use of publicly available big unstranded data with the stranded data could not only enhance detection of new transcripts but also reduce the generation of erroneous transcript models.

RNA-seq-based transcriptome assembly is also challenged by the imprecise ends of assembled transcripts (Steijger et al. 2013). Early methods roughly defined the transcription structures with the support of histone modification signals, such as H3K4me3 for active promoters and H3K36me3 for active gene bodies (Guttman et al. 2010; Ulitsky et al. 2011). Later, specialized RNA sequencing techniques, such as cap analysis gene expression by sequencing (CAGE-seq) (Yamashita et al. 2011; Brown et al. 2014; Kawaji et al. 2014) and poly(A) position profiling followed by sequencing (3P-seq) (Ulitsky et al. 2011; Nam and Bartel 2012), have been successfully applied to define the ends of transcripts at single base resolution. Integrative analysis of the specialized RNA-seq data including CAGE-seq and poly(A)-seq enabled the identification of more complete gene structures (Boley et al. 2014), valuable information for functional studies of the genes. However, despite the precise boundary information, the data are produced in a limited number of cell types. Recently, a computational method, GETUTR, that estimates 3’ UTR ends from general RNA-seq data was introduced (Kim et al. 2015), and it is now possible to predict the 3’ end of transcripts more accurately in any cell type using RNA-seq data.

Here, we developed a high-performance pipeline for transcriptome assembly, called the Co-Assembly of stranded and unstranded RNA-seq data Followed by End-correction (CAFE). Using the pipeline, the directions of a total of eighty-nine billion unstranded RNA-seq reads from multiple sources were successfully predicted and the reads with predicted directions (RPDs) were used for transcriptome assembly with stranded reads. The co-assembly of the RPDs and cell type-matched or pooled stranded RNA-seq reads improved the original assemblies of unstranded RNA-seq reads, and reconstructed a larger set of full-length transcripts. Of the resulting transcripts, thousands of putative lncRNAs were newly identified in addition to tens of thousands of known lncRNAs.

## RESULTS

### Unstranded RNA-seq causes error-prone assembly

To investigate the inaccuracy of transcriptome assemblies reconstructed from unstranded RNA-seqs (unstranded assemblies) relative to that of assemblies from stranded RNA-seqs (stranded assemblies), we reconstructed 45 stranded and 32 unstranded assemblies from publicly available RNA-seq data from the ENCODE project using Cufflinks (Trapnell et al. 2010). The resulting assemblies were evaluated based on the protein-coding genes of the GENCODE V19 annotations at the base level using Cuffcompare (Trapnell et al. 2010). The evaluation was done by counting false negative (FN), false positive (FP), and true positive (TP) bases upon agreement between the reference and the resulting assembly at the base level (Figure S1A; see Methods for more details). The sensitivity (TP/(TP+FN)) of the resulting assemblies appeared to be simply correlated with the size of mapped reads up to about 100 million mapped reads but converged beyond that size (Figure 1A), which suggests that many samples from ENCODE still need more data to reach their maximum sensitivity. On the other hand, the specificity (TP/(TP+FP)) of the unstranded assemblies was much less than that of the stranded assemblies when the resulting assemblies were evaluated with their directionality considered, regardless of the size of the mapped reads (Figure 1B). This result indicates that stranded reads provide more accurate information for transcriptome assembly. Previous studies failed to recognize the low accuracy of unstranded assemblies because they used a default option of Cuffcompare that ignores the directionality of the resulting assemblies during evaluation. In fact, the overall specificity, when the directionality of the resulting assemblies is not considered, neither correlates with the size of mapped reads nor differs between stranded and unstranded assemblies (Figures S1B and C).

**Figure 1.**
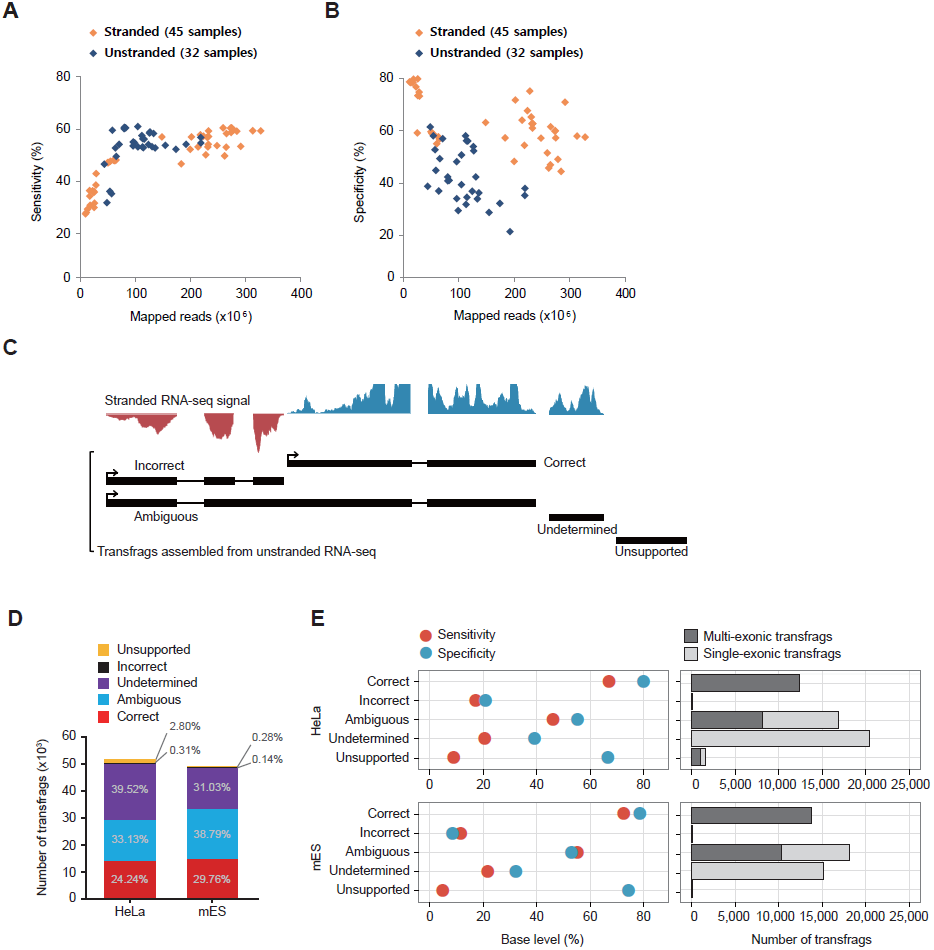
**Error-prone unstranded transcriptome assembly** (A-B) Sensitivities (A) and specificities (B) of stranded (orange diamond) and unstranded (navy diamond) assemblies constructed from ENCODE RNA-seq data are shown over the number of mapped reads. (C) Classification of transfrags assembled from unstranded RNA-seq data. Graphs on the top are signals from stranded RNA-seq data (blue is the signal in the forward direction and red is the signal in the reverse direction). Transfrags were classified into five groups according to the concordance and presence of the stranded signal: i) the correct group includes transfrags with more than 99% stranded reads in the same direction, ii) the incorrect group contains transfrags with more than 99% stranded reads in the opposite direction, iii) the ambiguous group contains transfrags with at least 1% and at most 99% stranded reads (at least three reads) in the opposite direction, iv) the undetermined group includes transfrags, all of which are single-exon, whose direction was not determined, v) the unsupported group includes transfrags with no stranded signal support. The 99% criterion was determined by systematic analyses with varying percentages (Figure S2). (D) Shown are the percentages of transfrags belong to the five groups - correct (red), ambiguous (blue), undetermined (purple), incorrect (black), and unsupported (yellow) in HeLa and mES cells. (E) The specificity (light blue) and sensitivity (red) of the five groups compared to the reference protein-coding genes in HeLa (left top) and mES cells (left bottom). The number of multi-exonic (dark gray) and single-exonic (gray) transfrags are indicated in each group (right).

To examine the nature and cause of the errors in unstranded assembly, we next sequenced both stranded and unstranded RNA-seq libraries that were simultaneously prepared in mouse embryonic stem (mES) cells. We also obtained a pair of publicly available stranded and unstranded RNA-seq datasets from human HeLa cells from the NCBI gene expression omnibus (GEO). These reads were mapped to reference genomes (hg19 for human and mm9 for mouse) using TopHat (Table S1), and unstranded reads (~68 million mapped reads for mES cells and ~40 million mapped reads for HeLa cells) were assembled using Cufflinks (Table S1). In total, 51,045 and 48,509 transcript fragments (transfrags) whose full lengths were not examined were assembled from HeLa and mES cells, respectively (Table S2). The resulting transfrags were divided into five groups based on their directions validated by stranded RNA-seq signals: correct, incorrect (those with an RNA-seq signal on the opposite strand), ambiguous (those with RNA-seq signals on both strands), undetermined (those with no direction), and unsupported (those with no stranded RNA-seq signals in either direction) (Figure 1C). All transfrags in the correct group (24.24% for HeLa cells and 29.76% for mES cells) were multi-exonic (Figures 1D and E); this high accuracy was the result of exon-junction reads that define the direction of the resulting intron with the splice-signal ‘GU-AG’ at the ends of the intron (Figure 1E). The remainder were regarded as problematic transfrags (75.76% for HeLa cells and 70.24% for mES cells). They displayed low accuracies and were placed in the incorrect (0.31% and 0.14%), ambiguous (33.13% and 38.79%), undetermined (39.52% and 31.03%), and unsupported (2.8% and 0.28%) groups (Figures 1D and E). They appeared to be severely defective in their structure and/or direction (Figure S3), and the majority in the undetermined group were single-exonic transfrags (Figure 1E). However, except for those in the unsupported group (Figure 1C), the defective transfrags (72.96% for HeLa cells and 69.96% for mES cells) could be corrected using the guide of the matched, stranded RNA-seqs.

### Probabilistic estimation of the directions of unstranded RNA-seq reads

To facilitate stranded assemblies with additional stranded reads, we sought to predict the directions of unstranded RNA-seq reads using *k*-order Markov chain models (*k*MC) whose transition probabilities were estimated with the directions of a current read x and its *k*-nearest stranded reads, x_*k*_. In the prediction step, the direction of a read with an unknown direction, y, was determined using maximum likelihood estimation (MLE) (Figure 2A). A read with a predicted direction (RPD) was treated as a stranded read and was used in the downstream assembly. For unstranded paired-end reads, the direction of a fragment was independently predicted. If the predicted direction of a fragment was not consistent with that of another fragment, the direction with a greater probability was chosen. Merely less than 1% (0.28% for HeLa and 0.14% for mES) were paired-end reads with discordantly predicted strands (Figure S4). Performing systematic analyses while increasing *k*, we found the optimum to be *k*=3, a value at which the accuracy is maximized and the computational cost is minimized (Figures S5A and B). Compared to a simple majority voting method with *k-*nearest stranded reads, *k*MC performed better as *k* increased (Figures S5C and 5D). Thus, we predicted the directions of all unstranded RNA-seq data using the Markov chain model with the optimum *k*-order and assembled all stranded read-like RPDs. Compared to the original assembly (unstranded assembly), those that were re-assembled from RPDs (RPD assembly) were significantly improved by 9.3–10.7% in their specificity without compromising their sensitivity (Figure 2B for HeLa cells and Figure 2C for mES cells). For instance, unstranded reads from a genomic locus where LOC148413 and MRPL20 are convergently transcribed were assembled into an erroneous annotation but their RPDs led to correction of the erroneous gene structure (Figure 2D). The prediction of strand information also significantly improved the specificity of the annotations of antisense-overlapped loci without compromising the sensitivity at the base level (Figures S6A and B).

**Figure 2.**
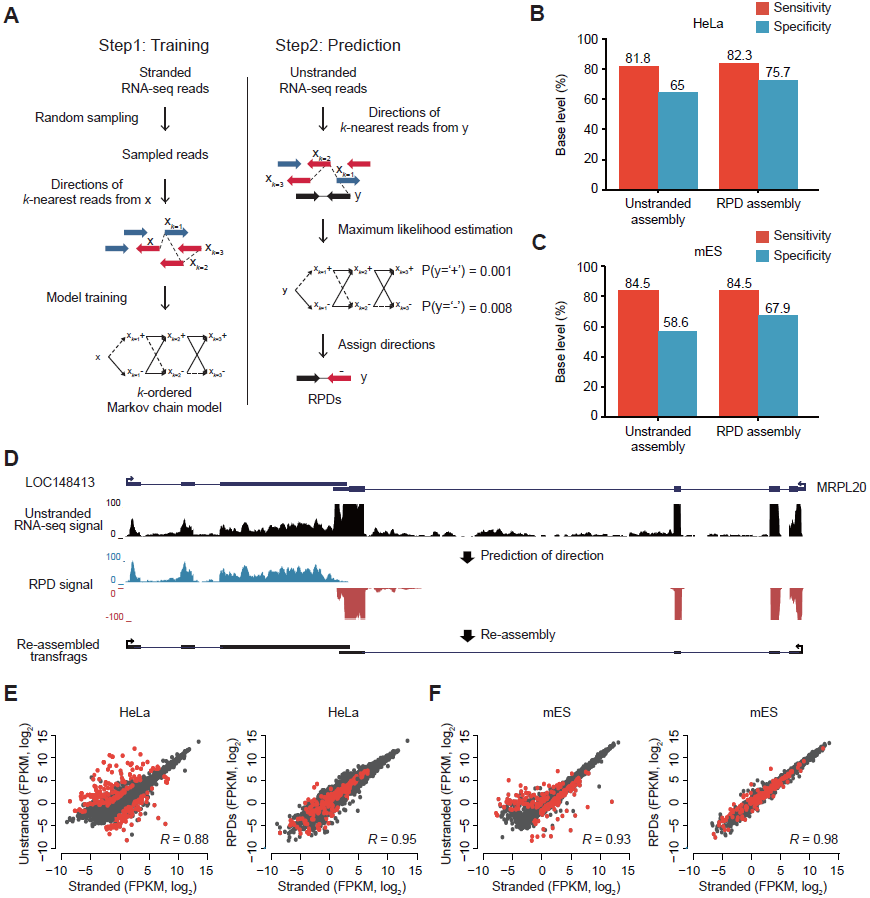
**Prediction of read directions using MLE** (A) Overview of *k*MC training and MLE of read direction. (Left) *S* base reads randomly sampled from stranded RNA-seq reads and their matched step-wise *k*-nearest reads (x_*k=1*_,x_*k=2*_,x_*k=3*_,…) were used for training *k*MC. Blue arrows are reads in the forward (+) direction and red arrows are reads in the reverse (-) direction. (Right) Prediction of read direction using MLE. Step-wise *k*-nearest stranded reads (x_*k=1*_,x_*k=2*_,x_*k=3*_,…) from a query unstranded read (black arrow) were extracted, and used to calculate two likelihoods at (+) and (-). A direction with the maximum likelihood is finally assigned to the query read. (B-C) Accuracies of transcriptomes assembled with RPDs (*k*=3) and unstranded reads in HeLa (B) and mEC cells (C). (D) An example of resulting transfrags re-assembled with RPDs. LOC148413 and MRPL20 are convergently overlapped at a locus where unstranded RNA-seq signals(black) are not separated but blue and red RPD signals are clearly separated in the forward and reverse directions, respectively. (E-F). Comparisons of gene expression values (FPKM, log_2_) estimated by stranded (X-axis) and unstranded reads (Y-axis, left) or RPDs (Y-axis, right) in HeLa (E) and mES cells (F). The correlation coefficients were calculated with Pearson’s correlation between the X- and Y-axis values. The red dots indicate genes with antisense-overlapped genes.

**Figure 3.**
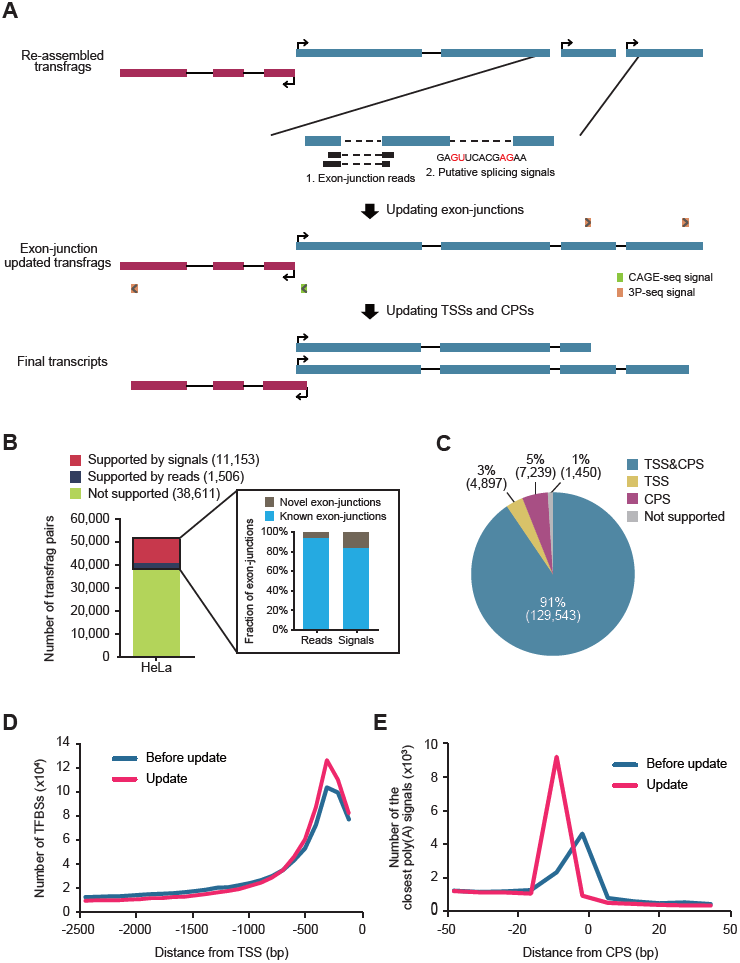
**Updating exon-junctions, TPSs, and CPSs in transfrag models** (A) Shown is a workflow for updating transfrag models, which comprises two steps: i) updating exon junctions and ii) updating TSSs and CPSs. (B) The number of neighboring transfrag pairs supported by putative splicing signals (red), by exon-junction reads (navy), and by neither (olive) in HeLa cells. The numbers in parentheses in the key indicate the number of pairs in each group. Among exon junctions supported by either exon-junction reads or putative splicing signals, the fractions of known (cyan) and novel (gray) exon junctions in GENCODE annotations are shown in the inset. (C) The fraction of transfrags updated with both TSS and CPS (blue), with only TSS (yellow), with only CPS (magenta), and with neither TSS or CPS (grey) in HeLa cells. (D) The number of TFBSs upstream of the original 5’ end (blue) and of the end updated with a TSS (pink) in HeLa cells. (E) The number of transfrags with a close poly(A) signal, AAUAAA, over the relative distances from the original 3’ end (blue) and the 3’ end updated with a CPS (pink) of transfrags in HeLa cells.

**Figure 4.**
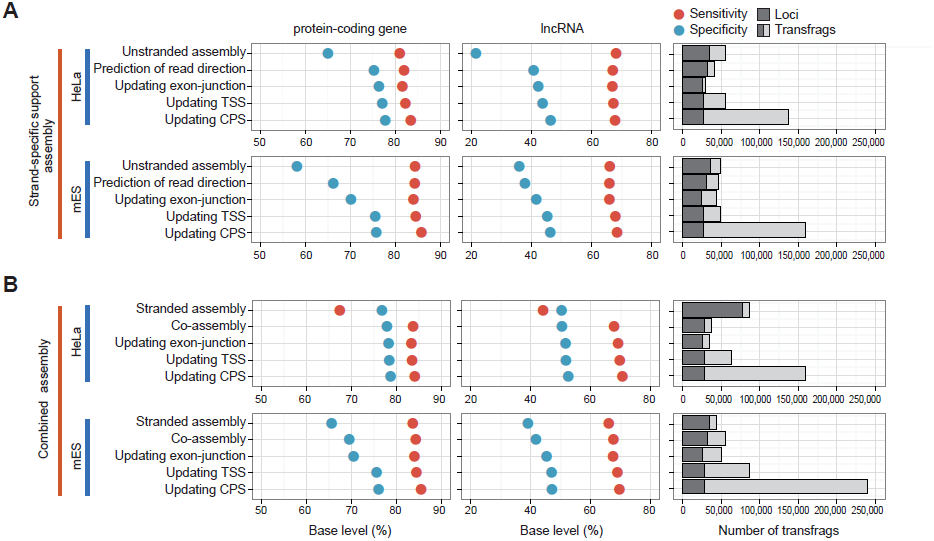
**Step-wise evaluation of transcriptomes re-assembled by CAFE** (A) Shown are the accuracies and sizes of strand-specific support transcriptomes (RPD assembly) at each step of CAFE in HeLa (top) and mES cells (bottom). The sensitivity (red solid circle) and specificity (blue) of the assemblies are measured by comparing to GENCODE protein-coding genes (left panel) and lncRNAs (middle panel). The number of assembled transfrags and their loci are indicated at each step (right panel). (B) Shown are the accuracies and sizes of combined transcriptome assemblies of both stranded reads and RPDs. The low sensitivity of the stranded assembly from HeLa cells is presumably because the stranded reads are of the single-end type and are 36 or 72 nt long. Otherwise, as in (A).

**Figure 5.**
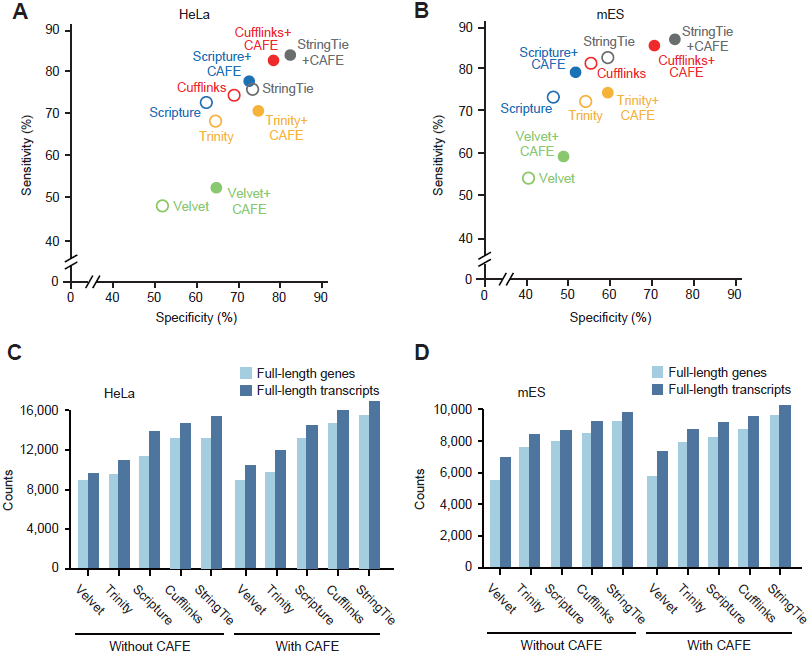
**Benchmarking other base assemblers** (A-B) The accuracies of combined transcriptome assemblies (solid circles) reconstructed by CAFE with base assemblers and of the original transcriptome assemblies (open circles) reconstructed by respective base assemblers, such as Cufflinks (red), Scripture (blue), StringTie (grey), Velvet (green), and Trinity (yellow), in HeLa (A) and mES cells (B). The accuracies of the original assemblies were calculated by averaging the accuracies of stranded and unstranded assemblies reconstructed by each base assembler. Velvet and Trinity were used as a *de novo* assembler, and Scripture, StringTie and Cufflinks were used as a reference-based assembler. (C-D) The numbers of full-length genes (light blue) and transcripts (blue) in the co-assemblies were compared to those in the original assemblies from HeLa (C) and mES cells (D). For the original assemblies, the higher number of full-length genes in the stranded and unstranded original assemblies was chosen.

**Figure 6.**
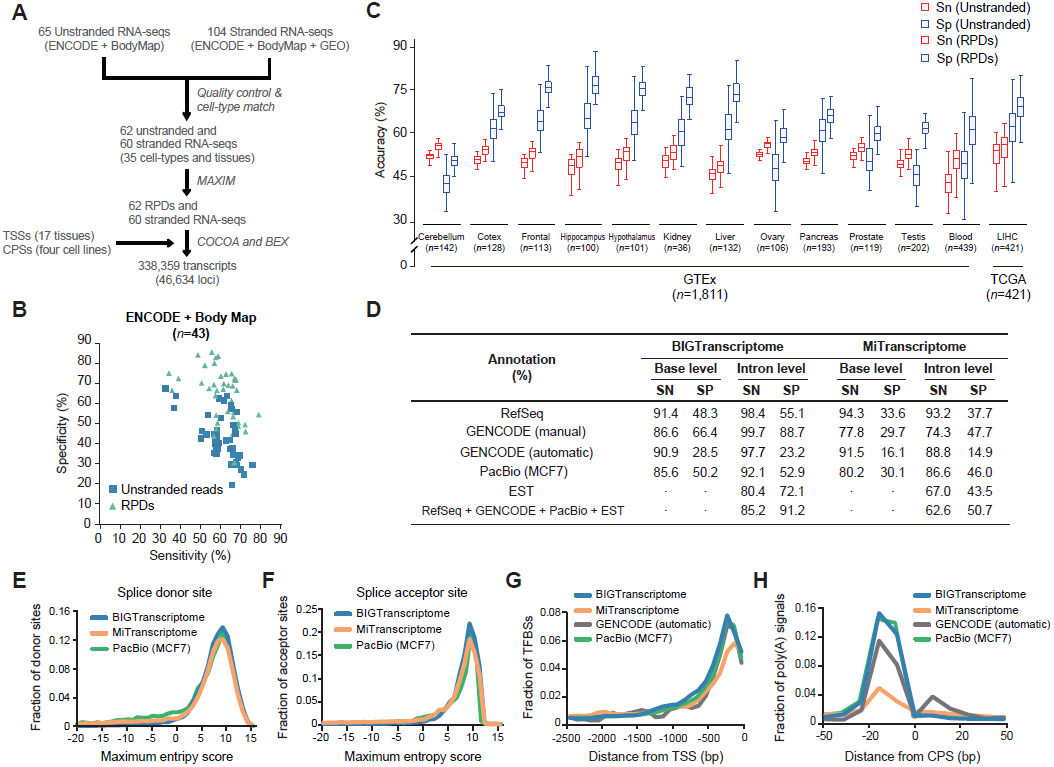
**Comprehensive human transcriptome map** (A) A schematic flow for the reconstruction of the BIGTranscriptome map using large-scale RNA-seq samples from human cell lines, ENCODE, and human BodyMap projects. (B) Accuracies of unstranded (blue) and RPD assemblies (mint) from the ENCODE and human BodyMap projects. (C) Sensitivities (red) and specificities (blue) of unstranded assemblies (solid line box) and RPD assemblies (dotted line box) are shown in box plots. The unstranded RNA-seq data are from GTEx (12 tissues) and the TCGA project (liver cancer). The numbers (*n*) indicate the sample numbers in each group. (D) Shown are the accuracies of BIGTranscriptome and MiTranscriptome at the base and intron levels based on four different sets of annotations (RefSeq, manual and automatic GENCODE, PacBio, and EST), and a combined set of annotations. SN: sensitivity and SP: specificity. (E-F) Maximum entropy scores of the putative splice donor sites (E) and of putative splice acceptor sites (F). Blue lines are from BIGTranscriptome, green lines are from PacBio assembly, and orange lines are from MiTranscriptome. (G) The fraction of TFBSs upstream of the 5’ end of BIGTranscriptome transcripts (blue) was compared to those of MiTranscriptome (orange), GENCODE (automatic) (black), and PacBio assembly (green). (H) The fraction of the closest poly(A) signals, AAUAAA, in the region just upstream of the 3’ end of BIGTranscriptome annotations (blue) compared to those of MiTransciprotme (orange), GENCODE (automatic) (black), and PacBio assembly (green).

The use of stranded RNA-seq data leads not only to better transcriptome assembly, but also in principle to better gene expression quantification. To test whether the expression quantification benefits from the prediction of strand information, the gene expression values were calculated with unstranded and corresponding RPDs, and then were compared to those calculated with stranded reads (Figures 2E-F and S6C). Overall, the unstranded reads over-estimated the expression level of genes in the loci with antisense-overlapping transcripts but RPDs corrected the over-estimation, leading to better correlation with those of stranded reads.

To test the general usage of the *k*MC model, we predicted the directions of unstranded reads from HeLa cells using the *k*MC model trained in mES cells, and vice versa. The species-mismatched models were comparable to the species-matched models (Figure S7), suggesting that the *k*MC model can be generalized to other cell types and species.

**Figure 7.**
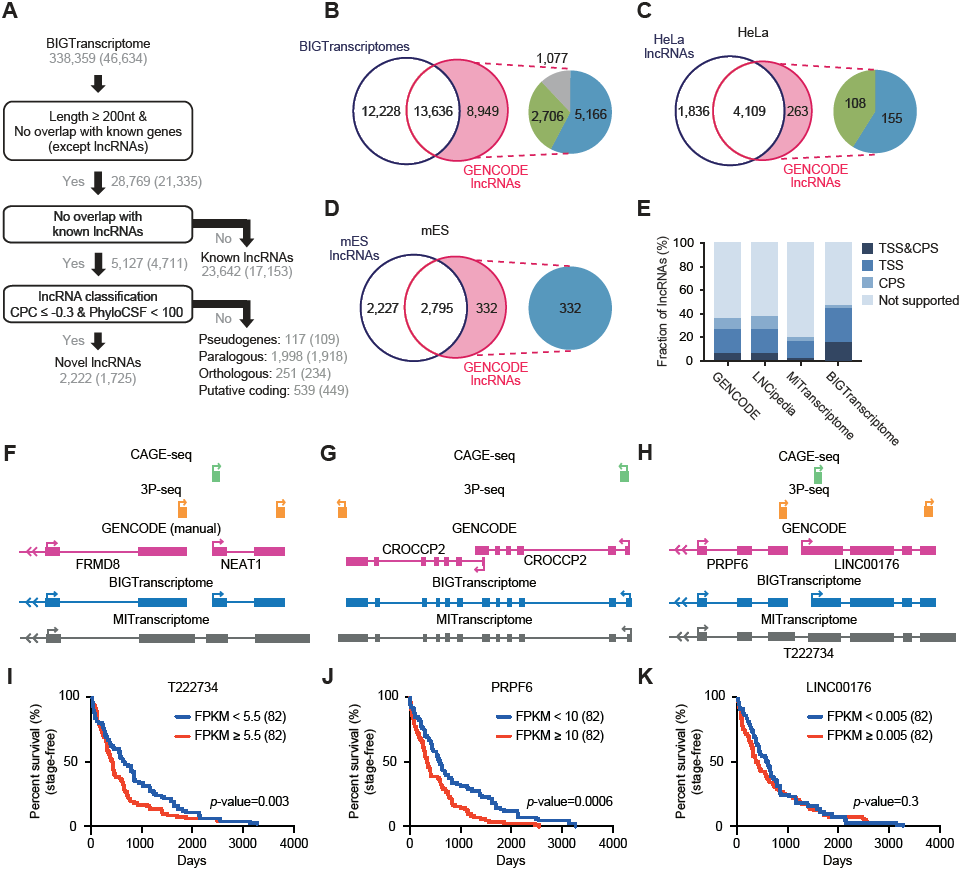
**BIGTranscriptome includes known and novel non-coding genes** (A) A schematic flow for annotating novel and known noncoding genes in BIGTranscriptome. (B) The Venn diagrams display the fraction of BIGTranscriptome lncRNAs that are published GENCODE lncRNAs. The inset indicates that GENCODE lncRNAs (8,949) not detected in BIGTranscriptome were classified as overlapping with known genes (blue), overlapping with falsely fused genes (green), or truly missed in our catalogue (grey). (C-D) Transcriptomes of HeLa (C) and mES cells (D) were compared to GENCODE lncRNAs, expressed over 1 FPKM in the matched cell types. The insets indicate that HeLa- and mES-expressed lncRNAs not detected in our lncRNA set were filtered by either overlap with known genes (blue) or mis-annotation (green). (E) The fractions of the indicated lncRNA sets with both TSS and CPS, either site, or neither site are shown in bar graphs. (F-H) Examples of mis-annotated gene models in public databases (MiTranscriptome and GENCODE). (F) The gene for a well-studied lncRNA, NEAT1, has been combined with a protein-coding gene, FRMD8, leading to mis-annotation as a protein-coding gene. (G) CROCCP2 is annotated in GENCODE (automatic) as having two independent isoforms whereas it is annotated as a single transcript in BIGTranscriptome and MiTranscriptome. (H) Gene models of BIGTranscriptome and MiTranscriptome, and CAGE-seq and 3P-seq data, at a locus. A fused single form, T222734, was annotated in MiTranscriptome whereas two independent genes, PRPF6 and LINC00176, were annotated in BIGTranscriptome. (I-K) Survival analyses for TCGA liver cancer samples based on the resulting gene models. 164 patient samples including termination events were divided into two groups, the top 50% (red) and bottom 50% (blue), by the median FPKM values of T222834 (I), PRPF6 (J), and LINC00176 (K).

### Refining boundaries and finding new exon junctions between transfrags

Shallow sequencing depth and short read length often cause transcript fragmentation in transcriptome assembly, mainly due to missing exon-junction reads and discontinuity of read overlaps. To improve the integrity of the assembled transcriptome, the missed exon junctions can be determined by either experimental (Clark et al. 2015) or computational approaches (Shapiro and Senapathy 1987; Reese et al. 1997; Pertea et al. 2001; Yeo and Burge 2004; Desmet et al. 2009). We first searched for potential exon junctions between neighboring transfrags on the same strand within a distance ranging from the first to the 99th percentile of the lengths of all known introns. After combining all exon-junction reads from all RNA-seq libraries, the potential junctions were examined and, if there were at least two junction reads that spanned two neighboring transfrags, the neighbors were connected (Figure 3A). Of 51,270 potential exon junctions, 1,506 (3%) were additionally supported by the method in HeLa cells (Figure 3B) and a similar fraction of potential junctions were supported in mES cells (Figure S8A). Of the newly connected exon junctions, 91.0–94.4% were present in GENCODE annotations (V19) and the remainder were novel (Figures 3B and S8A). The unconnected potential exon junctions were examined further with the program MaxEntScan (Yeo and Burge 2004) to determine whether the most likely putative splicing signal, ‘GU-AG,’ existed in the region between two neighboring transfrags (Figure 3A). Using that approach, 11,153 potential junctions for HeLa cells and 7,634 for mES cellswere newly connected (Figures 3B and S8A); 84.7–85.2% were present in GENCODE gene annotations and the remainder were novel (Figures 3B and S8A).

RNA-seq-based transcriptome assembly often results in imprecise transcript boundaries (Figure S9). To improve transfrag boundary annotation, transcription start sites (TSSs), determined from publicly available CAGE-seq (Consortium et al. 2014), and cleavage and polyadenylation sites (CPSs), determined from poly(A) position profiling by sequencing (3P-seq) (Nam et al. 2014), were incorporated into relevant transfrags. For TSSs and CPSs, respectively, 93–94% and 96–98% of transfrags were either confirmed or revised (Figures 3C and S8B). Transfrags updated for both TSS and CPS (91–92%) were regarded as full-length transcripts (Figures 3C and S8B). Updating TSSs improved the definition of the upstream promoter regions in which transcription factor binding sites (TFBSs) are significantly enriched (Figure 3D). Similarly, transfrags with CPSs displayed an enriched poly(A) signal, AAUAAA within 15–30nt upstream of the cleavage site, compared to those without CPS updates (Figure 3E).

### CAFE improves transcriptome annotations

We developed a pipeline, CAFE, which utilizes both stranded and unstranded RNA-seq data to reconstruct full-length transcripts effectively (Figure S10). To evaluate the pipeline, we first sought to re-assemble only RPDs (named strand-specific support assembly) from HeLa and mES cells, and measured the accuracy of intermediate assembly at each step by comparing our results to GENCODE protein-coding genes in the base level (Figure 4A). After updating TSSs and CPSs, the evaluation was proceeded with only transfrags with a major TSS and CPS while the count of transfrags took account of all isoforms. In total, 143,129 transfrags from 25,118 loci were assembled from HeLa cells; the quality of the resulting assembly for protein-coding genes was improved by about 14% for specificity and about 1.6% for sensitivity, compared to the original unstranded assembly (Figure 4A). Similarly, CAFE assembled 164,423 transfrags from 24,605 loci in mES cells and improved the quality of protein-coding gene assembly by 18.4% for specificity and 1.3% for sensitivity (Figure 4A). Although the resulting transfrags that overlapped with GENCODE lncRNAs were relatively less accurate than those of protein-coding genes partly because of their low and condition-specific expression patterns, CAFE also improved the quality of such transfrags by 22.1% and 8.3% for specificity in HeLa and mES cells, respectively, without compromising sensitivity. A major factor behind the increased specificity for both protein-coding and lncRNA genes was the prediction of read direction and re-assembly (Figure 4A).

We next performed combined assembly (co-assembly) of both stranded reads and RPDs using CAFE. The resulting assemblies included 166,227 transfrags from 25,591 loci in HeLa cells, of which 93.26% had their own TSSs and 94.62% had their own CPSs, and 244,085 transfrags from 26,332 loci in mES cells, of which 94.83% had their own TSSs and 98.08% had their own CPSs (Figures 4B and S11). Both the sensitivity and specificity of the final resulting transcriptome that overlapped with the GENCODE genes were greatly improved in the base level, compared to that in both of the original assemblies, and were slightly improved compared to the strand-specific support assembly (Figures 4 and S12).

### Benchmarking other transcriptome assemblers

To check whether the improvement in transcriptome assembly depends on a specific base assembler (originally, Cufflinks+CAFE), other reference-based assemblers, Scripture (Guttman et al. 2010) and StringTie (Pertea et al. 2015), were benchmarked using the same dataset (Scripture+CAFE and StringTie+CAFE). The resulting assemblies were more accurate for both HeLa (Figure 5A; 8.6~9.9% greater sensitivity and 11.4~12.9% greater specificity) and mES cells (Figure 5B; 3.2~4.9% greater sensitivity and 10.2~10.6% greater specificity) than the original assemblies in the base level. Because the recently published reference-based assembly pipeline GRITS utilizes only strand-specific paired-end reads, we excluded it from the benchmarking. Additionally, two available *de novo* assemblers, Trinity and Velvet, were also benchmarked by predicting the strand information of unstranded reads using CAFE and the resulting *de novo* assemblies of RPDs and stranded reads were more accurate than the original *de novo* assemblies (Figure 5A and B). Taken together, CAFE was able to improve initial assemblies robustly regardless of the base assembler used.

The number of full-length transcripts is another important aspect in the quality of transcriptome assembly. We thus compared the number of full-length transcripts assembled by CAFE to the number in the original and *de novo* assemblies. For these comparisons, transcripts that simultaneously included a TSS in the first exon and a CPS in the last exon were considered to be full-length transcripts. Trinity+CAFE and Velvet+CAFE assembled 8.8~10.4% more full-length transcripts than in the original *de novo* assemblies (Figure 5C). Cufflinks+CAFE, StringTie+CAFE, and Scripture+CAFE assembled 14.6%, 10.1%, and 13.9% more full-length transcripts than in the original assembly, respectively (Figure 5C). Similarly, CAFE constructed more full-length transcripts than in the original and *de novo* assemblies from mES cells (Figure 5D).

### High-confidence human transcriptome map

To construct a comprehensive human transcriptome map, large-scale transcriptome data were collected from the ENCODE project, the human BodyMap project, and GEO human cell lines; these data included 65 unstranded and 104 stranded RNA-seqs, TSS profiles across 17 human tissues, and CPS profiles from four human cell lines (Nam et al. 2014). We first predicted the directions of approximately six billion reads from 62 unstranded RNA-seq datasets using 60 cell-type-matched stranded RNA-seq datasets from 35 different cell types (Figure 6A and Table S3). The transcriptome assembly of the RPDs was more accurate than the unstranded transcriptome assembly in the base level (Figure 6B), suggesting that the prediction of read directions significantly reduced erroneously assembled transfrags. The co-assembly of RPDs and stranded reads with TSS and CPS profiles (Figure 6A) reconstructed 338,359 transcripts from 46,634 loci, named BIGTranscriptome. To expand our transcriptome map, we additionally predicted the strand information of > 2,200 individual unstranded RNA-seq data from 13 different tissues and tumors from the GTEx (Consortium 2013) and TCGA projects (Ciriello et al. 2013; Kandoth et al. 2013), and reconstructed more accurate transcriptome maps using the RPDs than using the unstranded reads at the base level (Figure 6C). The co-assembly of RPDs and stranded reads with TSS and CPS profiles using the sample pipeline shown in Figure 6A reconstructed tissue-specific transcriptome maps, named BIGTranscriptome-TS (Table S4). To examine their quality, all annotations were compared to those of RefSeq, GENCODE (manual), GENCODE (automatic), PacBio long read assembly (PacBio), and MiTranscriptome in terms of the number of full-length independent transcripts. Although BIGTranscriptome reconstructed fewer transcripts than did MiTranscriptome (Table S4), it contained more (16,376, 35.11%) independent genes that had at least one transcript with boundaries defined by TSSs and CPSs than did MiTranscriptome (5,741, 6.30%) and GENCODE (manual: 6,522, 13.59%; automatic: 1,301, 7.34%) (Table S5). Moreover, BIGTranscriptome included six thousand full-length independent single-exonic transcripts with a direction (~32.24% of single-exonic transcripts), whereas other annotations included tens of thousands of single-exon transcripts, only 1.04~20.91% of which were full-length independent single-exonic transcripts (Table S6A). Thousands of those that remained appeared to be partial fragments that were included in BIGTranscriptome annotations (Table S6B).

The accuracy of BIGTranscritpome annotations was also evaluated at the base level in terms of sensitivity and specificity based on RefSeq, GENCODE (manual), GENCODE (automatic), or PacBio (MCF7) annotations. BIGTranscriptome annotations were found to be 14.7 ~ 36.7% more specific for the RefSeq and GENCODE (manual) transcripts than were MiTranscriptome annotations, without compromising sensitivity (Figure 6D). We also checked if the intron structures of BIGTranscriptome agreed with those of the RefSeq, GENCODE, expression sequence tags (ESTs), PacBio, and combined annotations (RefSeq + GENCODE + EST + PacBio), and compared the results to those of MiTranscriptome. Overall, our BIGTranscriptome annotations were superior to those of MiTranscriptome for both sensitivity (22.6% greater) and specificity (40.5% greater) in the combined annotations (Figure 6D), indicating that BIGTranscritpome transcripts are less likely to be fragmented. 87.0% of the 29,274 putative BIGTranscriptome introns, not detected in the combined annotations, included a canonical splicing signal, ‘GU’-‘AG’, two nucleotides away from both ends; the remainder lacked the canonical splice signal (Table S7). Although the putative introns of MiTranscriptome also included the canonical splice signals at a similar level as BIGTranscriptome (Table S7), the putative splice sites of MiTranscriptome showed significantly lower maximum entropy scores than those of BIGTranscriptome at both splice donor and acceptor sites (Figures 6E and F). We also evaluated tissue-specific BIGTranscritome-TS annotations at the base and intron levels in terms of sensitivity and specificity based on RefSeq, GENCODE, EST, or PacBio annotations and found similar levels of accuracy in the transcriptome maps (Figure S13A).

To evaluate the accuracy of BIGTranscriptome transcript boundaries, we counted TFBSs in the regions upstream of the TSSs and canonical poly(A) signals in the regions around the CPSs. A higher fraction of TFBSs within 500nt upstream of a TSS (Figure 6G) and poly(A) signals within 15-30nt upstream of a CPS (Figure 6H) were observed for BIGTranscriptome transcripts than for MiTranscriptome and GENCODE (automatic), indicating that BIGTranscriptome includes transcripts with more precise ends. In addition, BIGTranscriptome agreed with PacBio and GENCODE about the 5’ and 3’-end positions of assembled transcripts better than did MiTranscriptome (Figures S13B-E). However, because the CPS information was profiled from only four human cell types, we additionally updated the cell-type specific 3’-ends of transcripts using GETTUR, which predicts the 3’-end of a transcript from RNA-seq data (Kim et al. 2015).

All RPDs of 62 unstranded RNA-seq samples, BIGTranscriptome, and BIGTranscriptome-TS annotations can be downloaded from our website (http://big.hanyang.ac.kr/CAFE).

### A confident catalogue of human lncRNAs

With our BIGTranscriptome map, we next sought to identify novel and known lncRNAs using the following lncRNA annotation pipeline, slightly modified from a previous method (Nam and Bartel 2012). Of 338,359 transcripts from 46,634 loci, 28,769 (8.5%) were longer than 200nt in length and did not overlap with exons of known protein-coding genes or ncRNA genes excepting lncRNAs. These transcripts were separated into known and putative lncRNAs. 23,642 transcripts from 17,153 genomic loci were previously annotated lncRNAs (Figure 7A). Putative lncRNAs were subsequently subjected to coding-potential classifiers: (1) coding potential calculation (CPC), which is a BLASTX-based similarity search method against all non-redundant protein sequences (Kong et al. 2007) and (2) PhyloCSF, which is a method to test whether an unknown sequence is evolved from coding or noncoding sequences using empirical codon models (Lin et al. 2011). Given the criteria for CPC (Figure S14) and PhyloCSF, 2,222 novel (from 1,725 loci) lncRNAs (Table S8) were identified as our human lncRNA catalogue (Figure 7A). The novel lncRNAs dominantly consisted of two exons and their length peaked at about 500 nt, similar to features of known lncRNAs (Figure S15). 2,905 transcripts (from 2,735 loci) that do not meet the lncRNA criteria were classified into pseudogenes (117 transcripts), paralogous transcripts (1,998), orthologous transcripts (251), and putative coding transcripts (539). Our known and novel lncRNA set can be downloaded from our website (http://big.hanyang.ac.kr/CAFE).

To evaluate our human lncRNA catalogues, genomic loci encoding GENCODE lncRNAs were compared to those of our catalogues. A majority (60.37% for GENCODE) were detected in our human lncRNA catalogues (Figure 7B). 7,872 (87.96%) of 8,949 GENCODE lncRNAs that were not in our catalogues were transcripts that overlapped with annotated genes in RefSeq, GENCODE, ENSEMBL, or MiTranscriptome (Figure 7B). We determined that 5,166 (65.63%) of 7,872 undetected GENCODE lncRNAs had been filtered out because they overlapped with known genes, and the remainder, 2,706, overlapped with a falsely fused transcript of a protein-coding gene and an lncRNA, mostly originating from MiTranscriptome (Figure 7B). For example, NEAT1, a well-studied lncRNA, is annotated as a protein-coding gene in MiTranscriptome because it was fused with FRMD8, an upstream protein-coding gene (Figure 7F). Only 4.76% (1,077/22,585) of GENCODE lncRNAs were truly missed in our catalogue. To verify this notion, the genomic loci encoding lncRNAs annotated from HeLa (Figure 7A) and mES cells (Figure S16) were respectively compared to those of GENCODE lncRNAs expressed at greater than 1 fragments per kilobase of exons per million mapped fragments (FPKM) in corresponding cells. We found that our HeLa and mES lncRNA sets included a majority of GENCODE lncRNAs (93.98% for HeLa and 89.38% for mES cells) (Figure 7C and D). All of the lncRNAs not included in our sets were transcripts that overlapped with known genes or that were mis-annotated in the public databases (Figure 7C and D). Similarly, the cell-type-specific lncRNA sets included more transcripts (~67% for HeLa and ~76% for mES cells) with fully or partially evident ends than did the non-cell-type-specific lncRNA catalogue (~48%) (Figure S17). Nevertheless, our lncRNA catalogue (13.45%) included many more intact gene models with fully evident ends than those of GENCODE, LNCipedia, and MiTranscriptome (6.61%, 6.71%, and 2.22%, respectively (Figure 7E). For example, CROCCP2 is shown to exist in two independent isoforms in GENCODE; however, it actually exists in a single form, as shown in both BIGTranscriptome and MiTranscriptome (Figure 7G).

We next sought to examine whether our BIGTranscriptome annotations could benefit the expression profiling of genes and their downstream analysis. T222734 was annotated as a single form in MiTranscriptome but this sequence turned out to be an independent protein-coding gene, PRPF6, and an lncRNA, LINC00176, evident with CAGE-seq and PolyA-seq, in BIGTranscriptome (Figure 7H). Using the single and the two independent forms of the genes, we performed survival analyses for 164 liver cancer samples from The Cancer Genome Atlas (TCGA). We found that the PRPF6 gene is a more significant marker (*p*-value: 0.0006) for the prognosis of the liver cancer patients than T222734 (*p*-value: 0.0034), whereas LINC00176 is expressed at a low level and is not significant marker (*p*-value: 0.31)(Figure 7I-K). Similarly, AC15645 (lncRNA) and MLXIP (protein-coding gene) were annotated in BIGTranscriptome but they were annotated as a single form (T087998) in MiTranscriptome (Figure S18A). The MLXIP annotated in our BIGTranscriptome appeared to be the more significant prognosis marker (Figure S18E) than T087998 and T088004 annotated in MiTranscriptome (Figure S18B and C) but the lncRNA, AC15645 turned out to be expressed at a low level (Figure S18D).

## Discussion

Our new transcriptome assembly pipeline, CAFE, enabled us to significantly improve the quality of the resulting assemblies by resurrecting large-scale unstranded RNA-seq data, which was formerly used for less informative or less specific transcriptome assembly. The re-use of the large-scale unstranded RNA-seq data could be valuable in three ways. For example, other public transcriptome databases, such as the TCGA consortium (Ciriello et al. 2013; Kandoth et al. 2013), the GTEx project (http://www.gtexportal.org) (Consortium 2013), the human protein atlas (Uhlen et al. 2010; Uhlen et al. 2015), and NCBI GEO, include large-scale unstranded RNA-seq data. Hence, determining the direction of unstranded sequence reads enables the construction of highly accurate transcriptome maps, which is necessary for highly qualitative downstream analyses. Although determining the direction of unstranded reads requires stranded data in the corresponding cell type or tissue, the use of pooled stranded data can still be of benefit to the prediction of transcript direction and the following assembly. In fact, the RPDs of unstranded Human BodyMap, GTEx, and TCGA data were predicted using pooled stranded RNA-seq data and showed better performance for specificity (Figures 6B and C). Secondly, in the case of genes with low expression such as those encoding lncRNAs, additional RPDs benefit transcriptome assembly by increasing the read-depth of those genes. Although the targeted capture of low-abundant transcripts like lncRNAs using antisense oligonucleotides enabled an increase in the copy number of the target transcripts (Clark et al. 2015), this approach is only applicable to known transcripts. Thirdly, additional RPDs could increase the detection of missed exon junctions, resulting in the connection of fragmented transfrags.

In this study, we utilized CAGE-seqs and 3P-seqs to profile transcript TSSs and CPSs, which detect unambiguous ends at single base resolution as well as transcript alternative forms. However, the assignment of multiple TSSs and CPSs raises a question: which pairs of ends, in all possible combinations, are relevant? Moreover, if a gene has alternative splicing isoforms, the number of possible isoforms is exponentially increased by multiple TSSs and CPSs. CAFE now generates all possible but unique isoforms, some of which would not actually exist in cells. Therefore, a precise way to determine a TSS-CPS pair simultaneously would provide biologically relevant isoforms directly. One approach is to integrate paired-end ditag (PET) data that contains both 5’- and 3’-end sequence tags of transcripts (Djebali et al. 2012) and an alternative is to sequence full-length RNAs using third-generation sequencing methods such as Iso-seq (Sharon et al. 2013).

Previous lncRNA catalogues suffered from transcript fragmentation, false extension and chimerization, and incorrect orientations of gene models. In fact, about 3 to 5% of documented lncRNAs are seemingly 3’UTR fragments of protein-coding genes. Moreover, because more than 9.5% of lncRNAs have a single exon, lncRNA gene direction should be re-examined. Our BIGTranscriptome map included many known and novel lncRNAs with unambiguous ends and more precise exon-intron models than previous maps. This comprehensive, precise transcriptome map should improve our understanding of the universe of noncoding genomes as well as the functional roles of noncoding RNAs.

## Methods

### Datasets

RNA-seq data used throughout this study were either from newly sequenced material or downloaded from public databases. A pair of stranded and unstranded RNA-seq datasets from mES cells were newly sequenced (GSE84946) and 169 publicly available samples of stranded and unstranded RNA-seq data from many different cell types, including HeLa and K562 cells, were downloaded from the ENCODE project (http://www.encodeproject.org"), human BodyMap project (http://www.broadinstitute.org), and our previous work (GSE52531). To collect data produced by a similar library construction method, samples were selected with the following criteria: 1) samples with poly-A selection applied; 2) samples with stranded RNA-seq data passing a quality control filter (such that the specificity of the stranded assembly is equal to or greater than 45%); 3) samples with unstranded RNA-seq and stranded RNA-seq data from the same cell type. After filtration, 122 RNA-seq samples across 35 cell types were analyzed for transcriptome assembly. In addition, 1,811 individual samples of unstranded RNA-seq data from 12 different cell types were downloaded from the GTEx project (http://www.gtexportal.org) and 421 hepatocellular carcinoma (LIHC) samples of unstranded RNA-seq were downloaded from TCGA project (Ciriello et al. 2013; Kandoth et al. 2013). The data were filtered with the same criteria above. 113 that did not meet with the criteria were excluded in the subsequent analyses. CAGE-seq processed data across 17 tissues for human and 23 tissues for mouse were downloaded from the FANTOM project (http://www.fantom.gsc.riken.jp). CPS data from many different cell types (HeLa, HEK293, Huh7, and IMR90 for human; mES, 3T3, liver, muscle, heart, white adipose tissue, and kidney for mouse) were downloaded from NCBI GEO (GSE52531).

### mES cell culture

mES cells were cultured in regular media containing 15% FBS, 1% PEN/STREP, 1% glutamine, 1% NEAA, and LIF. For mES cell maintenance, dishes were coated with 0.2% gelatin and irradiated CF1 mouse embryonic fibroblasts were plated as a confluent layer of feeder cells. mES cells were seeded at a density of 50,000 cells/6-well plate and were split every 2-3 days.

### RNA-seq library preparation

RNA from mES cells was extracted using an RNeasy kit (Qiagen, USA). Polyadenylated RNA was isolated using Oligo dT beads (Invitrogen, USA). Illumina Truseq stranded and unstranded mRNA library prep kits (Illumina, USA) were used for deep sequencing library preparation from 6 ug of total RNA according to the manufacturer's protocol. Libraries were sequenced in paired-end format to a length of 101bp using the HiSeq2000 platform (Macrogen Corporation, Republic of Korea).

### Preprocessing of RNA-seq data

To check RNA-seq data quality, RNA-seq reads were mapped to the corresponding reference genomes (hg19 for human and mm9 for mouse) using Bowtie (version 1.0.0) with default parameters. We calculated mismatch rates across the mapped read positions. If the raw read end(s) had a mismatch rate higher than 10%, they were trimmed off using Seqtk (version 1.0-r31). In addition, the trimmed reads with Phred base quality ≤ 20 were filtered using Sickle (version 1.200). The remaining reads were mapped to the corresponding reference genomes using Tophat (version 2.0.6) with mapping parameters “-i 61-l 265006 --min-segment-intron 61 --max-segment-intron 265006-g 5” for human and “-i 52-I 240764 --min-segment-intron 52 --max-segment-intron 240764-g 5” for mouse.

### Base transcriptome assembly

Base transcriptome assemblies were performed using Cufflinks (version 2.1.1) with assembly parameters “--min-isoform-fraction 0.15 --pre-mrna-fraction 0.2 --junc-alpha 0.001 --small-anchor-fraction 0.06 --min-frags-per-transfrag 12 -- max-multiread-fraction 0.65” for unstranded reads and “--min-isoform-fraction 0.05 --pre-mrna-fraction 0.2 --junc-alpha 0.01 --small-anchor-fraction 0.09 --min-frags-per-transfrag 8 -- max-multiread-fraction 0.65” for stranded reads.

### Benchmarking base assembly

To evaluate the performance of CAFE with other base assemblers, we performed reference-based assembly using Scripture (a beta version) with the default parameter and StringTie (version 1.3.0) with the parameter “-m 0-j 2-g 50-M 0.75” and *de novo* transcriptome assembly using Trinity (version 20140717) with the parameter “--min_contig_length 200” and Velvet (version 1.2.10) with the parameter “-hash_length 25-min_contig_lgth 50”. For benchmarking of reference-based assemblers (Scripture, Cufflinks, and StringTie) and *de novo* assemblers (Trinity and Velvet), each base assembler assembled transcriptomes using stranded and unstranded reads, respectively, and their averaged performance (sensitivity and specificity) was measured and then compared to the performance of CAFE in the co-assembly of RPDs and stranded reads.

### PacBio transcriptome assembly

To assemble the transcriptome from PacBio Iso-seq data, we downloaded data from human MCF7 cell lines sequenced with a total of 119 SMRT cells from the PacBio website (Biosciences). For subsequent analysis, we used the ‘Iso-seq’ protocol from the SMRT Portal provided by PacBio. Using the filtering module in the protocol, we acquired 1,857,590 reads of insert from Iso-seq data. In this step, we set parameters as “Minimum_Full_Passes 0 Minimum Predicted Accuracy 75”. Next, the classify module filtered about two-thirds of the reads of insert from the above with parameters set as “Minimum_Sequence_Length 300 Full-Length_Reads_Do_Not_Require_PolyA_Tails False”, leaving 524,084 full-length reads. The last module was the cluster module, which left 80,010 polished isoforms with “Predict_Consensus_Isoforms_Using_The_ICE_Algorithm True Call_Quiver_To_Polish_Consensus_Isoforms True Minimum_Quiver_Accuracy_To_Classify_ An_Isofrom_AS_HQ 0.99” parameters. Finally, GMAP (version 2015-07-23) was used with parameters “-f samse-n 0“ to map to the human genome, hg19. The final assembled set contained 47,416 non-redundant transcripts (15,688 genes).

### Reference gene annotations

Among the protein-coding genes and lncRNAs with “KNOWN” and “NOVEL” tags from GENCODE annotations (human: version 19, mouse: version M1), those with over 1 FPKM in the corresponding cells were selected. To build a bona fide lncRNA gene set, we performed the following filtration steps: (1) transcripts shorter than 200nt in length were discarded, and (2) lncRNAs sense-overlapping with exons of known protein-coding genes and noncoding genes (small ncRNAs including miRNAs, snRNAs, and snoRNAs and structural ncRNA genes including rRNAs and tRNAs) were excluded. As a result, 38,237 and 27,687 protein-coding genes and 4,380 and 1,328 lncRNAs were selected as reference gene annotations in human and mouse, respectively. The references were used to evaluate the quality of the transcriptome assemblies.

### **K-ordered Markov chain (*k*MC) models for read direction**

To predict the direction of unstranded reads mapped to the genome, *k*MC models were trained with the directions of the *k*-nearest stranded reads relative to a target read and with the direction of the target read. We built a training dataset including *S* base reads randomly selected from stranded reads mapped to genomes and their matched *k*-nearest reads. To acquire the *k*-nearest reads, we used a step-wise *k*-nearest method, in which the read x_*k=1*_ nearest to a query read x_*k=0*_ was first selected, then the read x_*k=2*_ nearest to the current read x_*k=1*_ was selected, then the read x_*k=3*_ nearest to the current read x_*k=2*_ was selected, and so on. To train unbiased models, we used 10 million as *S*, a large enough sampling number that is proportional to the *k* (also proportional to the number of states and edges to train). Practically, 2 X *K* matrix *M*_+_ or *M*_−_ for each emission value (+ and -) were constructed from the training data and each cell mor min the matrix indicates the fraction of + or − direction of the *j*th-nearest read x_*k=j*_ when the emission value (direction) of the previous state is *i*.

### Maximum likelihood estimation of read direction

The direction of an unstranded read, *r*, was inferred from the trained *k*MC models given step-wise *k*-nearest stranded reads of a query unstranded read mapped to a genome locus using MLE as in the following equation.

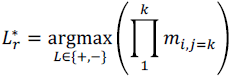

where *i* is the direction of the *k*th-nearest read, *L* is a set of possible directions that are {+, -}, and *L_r_^*^* is the maximum likelihood direction of the unstranded read *r*. Using the MLE, all maximum likelihood directions were predicted for all unstranded reads. If an unstranded read was paired-end, then its direction was determined differently, as follows. If a fragment of a paired-end read spanned an exon junction, the direction of the read was directly determined by the splice signal without MLE. If the directions of two fragments of a paired-read were inconsistent, the direction with greater likelihood was chosen for the read.

### Updating exon junctions

To update exon-junction signals missed in the original assembly, all pairs of neighboring transfrags within a distance ranging from 50 bp to the 99^th^ percentile of possible intron sizes (50–265,006 bp for human and 50–240,764 bp for mouse) were reexamined. The neighboring transfrags within a distance of 50 bp were combined using a default Cufflinks parameter, ‘--overlap-radius = 50’. If more than two exon-junction reads in at least two samples were detected, the neighbored transfrags were connected by the junction. Otherwise, the gaps between two neighboring transfrags were further scrutinized to detect *cis*-splicing signals. The gaps including splice donor ‘GU’ and acceptor ‘AG’ signals, but not TSSs or CPSs, between two neighboring transfrags were scanned by MaxEntScan (version 20040420), which calculates entropy scores for splice donor and acceptor sites. If the maximum entropy scores of both the splice donor and acceptor sites were above 0.217, a cutoff used in previous studies (Jian et al. 2014), then the interspace between the ‘GU’ and ‘AG’ was regarded as an intron and the two transfrags were connected by the intron.

### Updating TSSs and CPSs

The method for TSS identification from CAGE-seq tags was modified from the method for CPS identification from 3P-seq tags (Nam et al. 2014). Of the identified sites, those located in either the first exon or in the 3kb upstream region of a gene, without overlapping the upstream gene, were regarded as TSSs of the gene. Similarly, of the CPSs identified from 3P-seqs, those assigned to either the 3’UTR or the 5kb downstream region of a gene, without overlapping the downstream gene, were regarded as CPSs of the gene. After updating TSSs and CPSs, we removed all redundant transcripts or inclusive transfrags.

### Evaluation of transcriptome assembly

To evaluate the quality of transcriptome assembly, we compared the resulting assembly with the reference gene annotations (protein-coding genes and lncRNAs, respectively) using Cuffcompare (version 2.1.1). The sensitivity and specificity were estimated at the base and intron levels of the assembled transfrags. The base level sensitivity was calculated with the formula TP/(TP+FN), where TP is the count of nucleotides of the resulting transfrags falling within the reference protein-coding genes or lncRNA transcripts and FN is the count of nucleotides of the reference transcripts not falling within the resulting transfrags (Figure S1A). The base level specificity was calculated with the formula TP/(TP+FP), where FP is the count of nucleotides of the resulting transfrags not falling within the reference transcripts (Figure S1A). TP, FP, and FN were measured in a strand-specific manner. Transfrags without a determined direction are estimated on the assumption that there are either “+” and “-“ strands. When evaluating multiple isoforms, the union model was considered at the base level. However, for updating TSSs and CPSs, we chose the major isoform for the comparison. The intron-level sensitivity was also calculated with the formula TP/(TP+FN), where TP is the count of introns in the resulting transfrags that exactly match the introns (at both the 3’ and 5’ end) in the reference transcripts and FN is the count of introns in the reference transcripts that were not detected in the resulting transfrags (Figure S1A). The intron-level specificity was calculated with the formula TP/(TP+FP), where FP is the count of introns in the resulting transfrags that do not match the introns in the reference transcripts (Figure S1A). When evaluating multiple isoforms, the union set of introns were considered. To evaluate intron models, introns annotated in RefSeq, GENCODE, MiTranscriptome, and human ESTs from the UCSC genome database (https://genome.ucsc.edu/index.html) were extracted as references. Additionally, introns were compared to those detected via assembly of the long-read transcriptome from MCF7 cells. To acquire a confident set of introns from human ESTs, introns with a length within the range from the first to the 99^th^ percentile of all known introns and showing a canonical GU-AG splice signal were selected. For comparison, the exact match with the same direction and genomic position was used.

### Evaluation of full-length genes and isoforms

To evaluate how many full-length genes and isoforms were assembled, we collected the transcripts that simultaneously included a TSS in the first exon and a CPS in the last exon of the resulting transfrags. In addition, the transcripts aligned to the reference transcripts with at least a 95% match were regarded as full-length transcripts. At the gene level, gene models that unified all isoform exons were compared.

### Classification of lncRNAs

Of the putative lncRNAs, those with a CPC score ≤ -0.3 for human and ≤ -0.2 for mouse and a PhyloCSF score < 100 were classified as novel lncRNAs. Otherwise, they were classified as pseudogenes or orthologous, paralogous, or novel protein-coding genes. To find the optimal CPC score cutoff, we calculated CPC scores of reference protein-coding genes and lncRNAs in human and mouse, respectively. The cutoff at which the false discovery rate (FDR) was ≤ 0.01 for human and 0.05 for mouse and the true positive rate was maximized was chosen. The PhyloCSF score cutoff was derived from previous studies (Cabili et al. 2011; Kelley and Rinn 2012).

### Annotations of pseudogenes and paralogous, orthologous, and novel protein-coding genes

To annotate pseudogenes, we downloaded human and mouse pseudogene annotations from ENSEMBL (V74 for human and V60 for mouse). The transcripts overlapping pseudogene annotations were annotated as pseudogenes. Of the remainder, transcripts for which the longest open reading frames were significantly aligned (greater than 40%) to NCBI non-redundant (nr) database entries using BlastX were regarded as homologous genes. If the homologous gene was found in the corresponding species, it was regarded as a paralogous gene; otherwise, it was regarded as an orthologous gene. The genes with no significant homology were assigned as putative protein-coding genes.

### Survival analysis using TCGA liver cancer data

mRNA-seq BAM files and clinical information associated with LIHC samples were downloaded from the TCGA data portal (https://tcga-data.nci.nih.gov/). The expression levels of genes annotated in BIGTranscriptome, MiTranscriptome, and GENCODE were calculated using Cufflinks (Trapnell et al. 2010) as FPKMs. For 164 LIHC samples including termination events, survival analyses were conducted by Kaplan–Meier estimate (Dinse and Lagakos 1982). To examine whether the survival rates could be differentiated by expression levels calculated from the gene models of BIGTranscriptome, MiTranscriptome, and GENCODE, the samples were divided into two groups with the median FPKM value of the corresponding gene or transcript. The *p*-values were estimated using log-rank (Mantel-cox) test.

## Data accession

Sequence data can be accessed at the NCBI GEO using the accession number (GSE84946).

## Acknowledgements

This work was supported by the Basic Science Research Program through NRF, funded by the Ministry of Science, ICT & Future Planning (NRF-2013R1A1010185, NRF-2014M3C9A3063541, and NRF-2012M3A9D1054516).

## Supplementary Figure Legends

**Figure S1. Evaluation of assembled transfrags with or without consideration of their directionality** (A) Evaluation of assembled transfrags at the base and intron level. Resulting transfrags were compared to the reference gene structures and were counted as true positive (TP), false positive (FP), and false negative (FN) at the base (top) and intron level (bottom). Isoforms were combined to create a unified exon model for evaluation. When comparing to the reference in the forward strand, transfrags in the reverse strand were classified as FP when their directionality was considered but as TP when the directionality was not considered. See Methods for more details. (B-C) The sensitivity (B) and specificity (C) of the resulting assemblies produced from the ENCODE dataset. The Cuffcompare analysis was done without considering their directionality during evaluation (default option). The stranded (orange diamond) and unstranded (navy diamond) assemblies were separately evaluated.

**Figure S2. Criteria for transfrag classification** (A-B) Shown are the specificities of correct transfrags across strand rates and antisense read numbers. The strand rate and the antisense read number that display the maximum specificity were used as criteria (dotted box) for the classification of transfrags in HeLa (A) and in mES cells (B).

**Figure S3. Problematic transfrags assembled from unstranded RNA-seq** (A-C) Examples of problematic transfrags belong to three groups – ambiguous (A), undetermined (B), and incorrect (C). Graphs are signals from unstranded RNA-seq data.

**Figure S4. Concordant and discordant predictions of paired-end reads** (A-B) Percentage of concordant (blue), discordant (red), and exon-junction-support (orange) predictions of strand information for paired-end reads in HeLa cells (A) and mES cells (B).

**Figure S5. Systematic analysis of MC *k*-order** (A-B) Accuracies of RPDs across different *k*-orders. The optimal *k*-order (dotted box) was set as the *k* at which the specificity was maximized for HeLa (A) and mES cells (B). (C-D) Accuracies (F-scores and sensitivity) of RPDs by *k*MCs and *k*-nearest majority voting across different *k* values (from 1 to 10)for the antisense-overlapping loci in HeLa (C) and mES cells (D). F-score was calculated by the formula, 
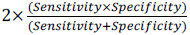

**Figure S6. Effectiveness of the *k*MC model** (A-B) Sensitivity (SN, red) and specificity (SP, blue) were measured for the resulting assembly across the antisense-overlapping loci with different fractions of antisense reads using unstranded reads (open circles) and RPDs (solid circles) from HeLa (A) and mES cells (B). (C) Comparisons of gene expression values (FPKM, log_2_) estimated with stranded (X-axis) and unstranded reads (Y-axis, left) or RPDs (Y-axis, right) from whole blood cells (Zhao et al. 2015). The gene expression values were averaged from four biological replicates of whole blood cells. Otherwise, as in Figure 2E-F.

**Figure S7. Generality of the *k*MC model** (A-B) Comparisons of unstranded, species-mismatched, and species-matched models. Accuracies of unstranded, species-mismatched (by the *k*MC models trained in mES cells), and species-matched models (by the *k*MC models trained in HeLa cells) in HeLa cells (A) and vice versa (B).

**Figure S8. Updating mES cell transcript models** (A) The number of neighboring transfrag pairs supported by putative splicing signals (red), by exon-junction reads (navy), and by neither (olive) in mES cells. Otherwise, as in Figure 3B. (B) The fraction of transfrags updated with both TSS and CPS (blue), with only TSS (yellow), with only CPS (magenta), and with neither TSS or CPS (grey) in mES cells.

**Figure S9. Erroneous transfrag boundaries** (A) Shown is an example of mis-assembly of PNPLA2 at its 5’ end, evident by CAGE-seq signals. (B) Shown is an example of mis-assembly of UBE2J2 at its 3’ end, evident by 3P-seq signals.

**Figure S10. A schematic flow of the CAFE pipeline** Shown is the schematic flow of the CAFE pipeline according to the combined, strand-specific support and strand-specific assembly. If there are both stranded and unstranded reads in the same cell type, the MAXIM, COCOA, and BEX steps are all executed. If there are only unstranded reads, the MAXIM step is carried out with the pooled stranded RNA-seq data.

**Figure S11. The fraction of transcriptomes co-assembled with RPDs and stranded reads** (A-B) The fraction of transcriptomes updated with both TSS and CPS (blue), with only TSS (yellow), with only CPS (magenta), and with neither TSS or CPS (grey) in HeLa (A) and in mES cells (B).

**Figure S12. Evaluation of transcriptomes assembled with only stranded reads** (A) Shown are the accuracies and sizes of strand-specific transcriptomes assembled using only stranded reads in HeLa (top) and mES cells (bottom). Otherwise, as in Figure 4.

**Figure S13. Accuracies of BIGTranscriptome-TS and boundary comparisons of BIGTranscriptome** (A) Shown are the accuracies of BIGTranscriptome-TS (13 tissues and cancer) at the base and intron levels compared with those of four different sets of annotations (RefSeq, manual and automatic GENCODE, PacBio, and EST) and a combined set of annotations. SN: sensitivity and SP: specificity. (B-C) Shown are the relative distances from the ends of GENCODE transcripts with the same splicing pattern to the ends of BIGTranscriptome isoforms (blue) versus to the ends of MiTranscriptome isoforms (orange). (B) for the 5’ end and (C) for the 3’ end. (D-E) Shown are the relative distances from the ends of the PacBio (MCF7) transcripts with the same splicing pattern to the ends of BIGTranscriptome isoforms versus to the ends of MiTranscriptome (orange). (D) for the 5’ end and (E) for the 3’ end.

**Figure S14. The optimal cutoff value of the coding potential calculator (CPC)** (A-B) Shown are the density graphs across different CPC scores of multi-exonic protein-coding genes (green), single-exonic protein-coding genes (purple), 3’ UTRs (sky), and Goldstandard lncRNAs (orange) in HeLa (A) and in mES cells (B).

**Figure S15**. **Basic statistics of known and novel lncRNAs** (A-D) The number of exons (A and C) and lengths (B and D) of known (A and B) and novel (C and D) lncRNAs in BIGTranscriptome. Vertical dotted lines indicate the medium length of lncRNAs.

**Figure S16. Annotations of novel and known lncRNAs in mES cells** A schematic flow for the annotation of novel and known lncRNAs in mES cells.

**Figure S17. The fraction of lncRNAs with evident TSS and CPS** (A-C) The fractions of lncRNAs with both TSS and CPS, either site, or neither site are shown in pie charts for BIGtranscriptome (A), HeLa cells (B) and mES cells (C). The numbers in parentheses indicate the number of lncRNAs that belong to each group.

**Figure S18. Survival analysis for liver cancer samples based on mis-annotated transcripts** (A) Gene models of a protein-coding gene, MLXIP, and a lncRNA, AC156455, in BIGTranscriptome and MiTranscriptome. (B-E) Survival analyses for TCGA liver cancer patient samples based on the gene models. 164 samples including termination events were divided into two groups, the top 50% and bottom 50%, according to the median FPKM values of T087998 (B), T088004 (C), AC156455 (D), and MLXIP (E).

